# Peak gamma frequency and cortical laminar processing are modified across the healthy menstrual cycle

**DOI:** 10.1101/219196

**Authors:** R.L. Sumner, R.L. McMilllan, A. D. Shaw, K.D. Singh, F. Sundram, S.D. Muthukumaraswamy

**Affiliations:** School of Psychology, The University of Auckland, Auckland, 1142, New Zealand; School of Pharmacy, The University of Auckland, Auckland, 1142, New Zealand; CUBRIC, School of Psychology, Cardiff University, Cardiff, CF24 4HQ, UK; Department of Psychological Medicine, The University of Auckland, Auckland, 1142, New Zealand

**Keywords:** Menstrual cycle, visual gamma oscillations, electroencephalography (EEG), GABA, allopregnanolone

## Abstract

Fluctuations in gonadal hormones over the course of the menstrual cycle are known to cause functional brain changes and are thought to modulate changes in the balance of cortical excitation and inhibition. Animal research has shown this occurs primarily via the major metabolite of progesterone, allopregnanolone, and its action as a positive allosteric modulator of the GABA_A_ receptor. Our study used EEG to record gamma oscillations induced in the visual cortex using stationary and moving gratings. Recordings took place during twenty females’ mid-luteal phase when progesterone and oestradiol are highest, and early follicular phase when progesterone and oestradiol are lowest. Significantly higher (~5 Hz) gamma frequency was recorded during the luteal compared to the follicular phase for both stimuli types. Using dynamic causal modelling these changes were linked to stronger self-inhibition of superficial pyramidal cells in the luteal compared to the follicular phase. In addition the connection from inhibitory interneurons to deep pyramidal cells was found to be stronger in the follicular compared to the luteal phase. These findings show that complex functional changes in synaptic microcircuitry occur across the menstrual cycle and that menstrual cycle phase should be taken into consideration when including female participants in research into gamma-band oscillations.

## Introduction

In healthy women, fluctuations in gonadal hormones lead to functional changes in the brain over the course of each menstrual cycle. The interaction of these hormones with the balance of cortical excitation and inhibition has formed the basis of research attempting to characterise changes across both the healthy menstrual cycle as well as menstrual cycle linked disorders, such as catamenial epilepsy and premenstrual dysphoric disorder (PMDD) (Bäckström et al., 2014; Reddy, 2004). As the primary mediator of cortical inhibition, the γ-aminobutyric acid (GABA) system and aberrant GABAergic inhibition has been implicated in these menstrual cycle related disorders (Bäckström et al., 2003). Over the course of the menstrual cycle, changes in the GABA system are modulated primarily by the major metabolite of progesterone, allopregnanolone. Allopregnanolone, similar to benzodiazepines, produces this change via its action as a potent positive allosteric modulator of the GABA_A_ receptor. By binding to the neurosteroidal site on the GABA_A_ receptor allopregnanolone potentiates the effect of GABA, leading to an overall increase in inhibition of neuronal excitability (Birzniece et al., 2006; Majewska, Harrison, Schwartz, Barker, & Paul, 1986). While the effect of allopregnanolone on the balance of cortical excitation and inhibition is beginning to become apparent in animal research (Smith, Shen, Gong, & Zhou, 2007), limitations in measuring functional changes *in vivo* in humans has meant a major gap in understanding remains.

Studies using Magnetic Resonance Spectroscopy (MRS) have found variations in levels of GABA both across the healthy menstrual cycle and in atypical populations. The first of these studies reported finding reduced GABA levels across the menstrual cycle in PMDD compared to healthy controls (Epperson, Haga, Mason, & et al., 2002). Furthermore, Epperson, Haga, Mason, Sellers, et al. (2002) found that GABA levels overall were increased in the luteal phase compared to the follicular phase in healthy controls. The opposite was found for participants with PMDD. Similar trends were found in a study that compared smokers with healthy controls (Epperson et al., 2005) indicating this effect was not specific to either group. However, GABA concentration, as measured with MRS, only provides a bulk concentration measurement and it is unclear how these measures directly index functional synaptic inhibition (Stagg, Bachtiar, & Johansen-Berg, 2011). However, these studies (Epperson, Haga, Mason, & et al., 2002; Epperson et al., 2005) do give evidence for significant functional changes in the human female GABA system over the course of the menstrual cycle.

A number of studies have also attempted to elucidate functional changes in the GABA system that occur over the menstrual cycle. Neurosteroids have been linked to changes in tonic inhibition as well as changes in receptor density (Lovick, Griffiths, Dunn, & Martin, 2005; Maguire, Stell, Rafizadeh, & Mody, 2005). Furthermore, direct evidence of allopregnanolone influencing changes in GABA_A_ receptor expression and sensitivity has been found in animals (Lovick et al., 2005; Maguire et al., 2005; Türkmen, Bäckström, Wahlström, Andreen, & Johansson, 2011). In humans, recent research using administered allopregnanolone indicated reduced sensitivity in the luteal phase compared to the follicular phase by showing reduced sedation in the luteal phase, measured using saccadic eye movement (SEM) (Timby et al., 2016). However, earlier research, also measuring SEMs, found the opposite using midazolam (a GABA enhancing drug) and administered pregnanolone (Sundstrom et al., 1998; Sundstrom, Nyberg, & Bäckström, 1997).

Neural oscillations recorded using EEG present another non-invasive method of studying humans and have been related to GABAergic inhibition. In particular beta oscillations have been linked to changes in GABAergic inhibition (Whittington, Traub, Kopell, Ermentrout, & Buhl, 2000). One study showed a relative decrease in beta power in the luteal phase (Solis-Ortiz, Ramos, Arce, Guevara, & Corsi-Cabrera, 1994). Increased beta power has been shown with a number of GABA enhancing drugs such as benzodiazepines (Feshchenko, Veselis, & Reinsel, 1997; Hall, Barnes, Furlong, Seri, & Hillebrand, 2010; Jensen et al., 2005; van Lier, Drinkenburg, van Eeten, & Coenen, 2004). Alpha oscillations have a well-known relationship with GABAergic inhibition (Jensen & Mazaheri, 2010). However, the mechanisms by which alpha is modulated by changes in GABA are less well established as the modulations that occur are not always predictable (Jensen & Mazaheri, 2010; Lozano-Soldevilla, ter Huurne, Cools, & Jensen, 2014). Alpha oscillations have also been correlated with changes over the menstrual cycle, though the results have been conflicting (Bazanova, Kondratenko, Kuzminova, Muravlyova, & Petrova, 2014; Brötzner, Klimesch, Doppelmayr, Zauner, & Kerschbaum, 2014; Solís-Ortiz, Ramos, Arce, Guevara, & Corsi-Cabrera, 1994).

Visually induced gamma oscillations are an attractive approach for understanding changes in cortical inhibition. Animal *in vivo* and *in vitro* studies have established that both power and frequency of gamma oscillations are intrinsically linked to changes in GABAergic inhibitory processes (Bartos, Vida, & Jonas, 2007; Gonzalez-Burgos & Lewis, 2008). Buzsaki and Wang (2012) illustrate how changes in synchronicity of gamma oscillations can be mediated by inhibitory GABAergic interneurons exerting their control on large populations of pyramidal cell firing. Via this model, increases in GABAergic inhibition leads to synchronisation of the previously stochastic firing of pyramidal cells. Electrophysiological recording demonstrates a decrease in frequency and an increase in gamma amplitude in response to increases in GABAergic inhibition (Buzsaki & Wang, 2012; Gonzalez-Burgos & Lewis, 2008). This is a useful model for both MEG and EEG recordings of gamma oscillations, as both methods are particularly sensitive to changes in local field potentials associated with cortical pyramidal cells (Buzsáki, Anastassiou, & Koch, 2012).

There is promising translation of this cellular model to non-invasive human studies via induction of gamma oscillations in the visual cortex. Visual gamma oscillations appear to be a highly robust trait biological trait marker (Muthukumaraswamy, Singh, Swettenham, & Jones, 2010; Tan, Gross, & Uhlhaas, 2016) that varies across age (Gaetz, Roberts, Singh, & Muthukumaraswamy, 2012; Orekhova et al., 2015), and is strongly genetically determined (van Pelt, Boomsma, & Fries, 2012). A number of pharmaco-MEG studies have been able to show that they are consistently and predictably modulated by changes in GABAergic inhibition (Muthukumaraswamy, 2014). Drugs that increase GABAergic inhibition via positive allosteric modulation including alcohol, and the benzodiazepine lorazepam have been shown to lead to decreases in gamma frequency and increases in gamma power (Campbell, Sumner, Singh, & Muthukumaraswamy, 2014; Lozano-Soldevilla et al., 2014; Magazzini et al., 2016). Similarly, tiagabine, a GABA reuptake inhibitor used in the treatment of epilepsy has been shown to decrease the frequency but not modify the amplitude of gamma oscillations (Magazzini et al., 2016). Furthermore, recent research has linked gamma oscillations in the human primary visual cortex (V1) to changes in GABA_A_ receptor density (Kujala et al., 2015).

Recently, dynamic causal modelling of steady-state responses (DCM-SSR) has been used to explain spectral data at the level of network changes within prescribed generative neural mass models. DCM allows inferences to be made about underlying microcircuitry of a cortical response acting as a “mathematical microscope” (Moran, Pinotsis, & Friston, 2013). Of particular interest in analyses of visually induced gamma oscillations is the ability to make inferences about intrinsic connectivity between populations of superficial pyramidal cells and inhibitory interneurons, potentially reflecting the neural dynamics underlying the generation of gamma oscillations (Buzsaki & Wang, 2012). Recently Shaw et al (2017) developed a customised canonical microcircuit model that better reflects the synaptic physiology of V1 than the generic mass models used in previous DCM work, in order to study visually induced gamma oscillations which arise from V1. Using their model of V1, Shaw et al. (2017) were able to show significant modulation of parameter strength when visually induced gamma activity was modified by tiagabine. Furthermore, they were able to differentiate between laminar specific events including localising the generation of gamma oscillations to primarily superficial layers through pyramidal-interneuron loops. The study was also able to show model sensitivity to the effect of tiagabine on the underlying time-constant of GABAergic inhibition. As such, this approach provides a potentially attractive method for understanding the changes in the GABA system across the menstrual cycle.

The aim of the current study was to quantify visual gamma oscillation parameters over the course of the menstrual cycle in healthy women. Furthermore, using the DCM approach developed by Shaw et al. (2017) we sought to make inferences about functional changes in the GABA system that may contribute to any changes found. The study used EEG to record visually induced gamma oscillations in the early follicular phase compared to the mid luteal phase, when hormones are at their lowest compared to their highest levels.

## Methods

### Participants and study design

Twenty females aged 21-23 years volunteered to participate in the study and completed both study days. Participants were required to have no history of neurological or psychiatric disorder including pre-menstrual dysphoric disorder or any self-reported major menstrual cycle related changes in mood. They were also required to not be taking any ongoing prescription medications or use hormonal forms of contraception. This study was approved by the University of Auckland Human Participants Ethics Committee. Participants provided informed written consent prior to participation.

The study used a two-session crossover design. One session took place during the follicular phase when oestradiol and progesterone are, on average, at their lowest point in the cycle while the other session took place in the mid- luteal phase when oestradiol and progesterone are, on average, at their peak in the menstrual cycle (Bäckström et al., 2003; Genazzani et al., 1998). The order of sessions was counterbalanced. All study sessions began between 2-4pm to control for diurnal variations in neurosteroid levels (Tiihonen Möller, Bäckström, Söndergaard, Kushnir, & Bergquist, 2016).

Participants were tracked for 3 menstrual cycles leading up to their first study date. This allowed the best estimate of cycle length to be made. Based on the average menstrual cycle length of 28 days, the follicular phase was defined as between day 1-5 of the menstrual cycle; the luteal phase, was defined as between 20-25 of the menstrual cycle. Several participants had considerably longer or shorter cycles than 28 days. For these participants, the luteal study date was adjusted accordingly. For this reason, blood samples were taken for confirmation of correct cycle timing.

### Blood samples

For participants with an average cycle length, blood samples were taken on the study day to test for progesterone and oestradiol levels in plasma. Participants with longer or shorter cycles came in for an additional sample the day before to confirm they were in the right phase. Cycle timing was determined using the guidelines provided by local laboratory guidelines (Lab Plus NZ), with progesterone levels used as the confirmation of timing. Blood samples were collected using BD vacutainer^®^ PST^™^ II tubes. Quantification of oestradiol and progesterone concentration in plasma was carried out by LabPlus, Auckland Hospital, by electrochemiluminescence immunoassay. Assays were performed according to Roche Oestradiol III (2016) and Progesterone III (2015) assay guidelines using a Roche Cobas 8000 analyser (e602 module).

### EEG acquisition and paradigm

Continuous EEG was recorded using 64 channel Acticap Ag/AgCl active shielded electrodes and Brain Products MRPlus amplifiers. Data were recorded in Brain Vision Recorder (Brain Products GmbH, Germany) with a 1000 Hz sampling rate, and 0.1 μV resolution. FCz was used as an online reference, AFz as ground. Electrode impedance below 10 kΩ was achieved prior to recording.

Stimuli were displayed on an ASUS VG248QE computer monitor with a screen resolution of 1920 x 1800 and 144 Hz refresh rate. TTL pulses generated through the parallel port of the display computer provided synchronisation of stimulus events with EEG acquisition. All stimuli were generated by MATLAB (The Mathworks, Inc., Natick, MA) using the Psychophysics Toolbox (Brainard, 1997; Kleiner et al., 2007; Pelli, 1997).

The stimuli for this task was a black and white annular grating with a spatial frequency of 3 cycles per degree, subtending 16 degrees of visual angle (Figure 1a). The grating was presented at 90% contrast, in the centre of the screen, on a grey background. A red dot provided a central fixation point. Participants were seated 90cm from the screen. There were two conditions in this experiment (Figure 1a). In condition one, the stimuli moved inwardly at a rate of 1.33 degrees of visual angle per second. In condition two, the stimuli were stationary. In both conditions the stimulus was on for between 1-1.1s (pseudorandomly jittered). Participants were instructed to press the spacebar as soon as the stimulus disappeared from the screen. If the response was too slow the paradigm paused while participants were prompted to “press space to continue”. Following a participant’s response there was then a 1s inter-trial interval. Each condition was presented 3 times each in a block of 84 presentations. The order of blocks was counterbalanced. Each block was separated by a forced 20s break. Two participants only received the moving grating condition, leaving 16 participants for the static gratings analyses and 18 for the moving grating analyses.

### Data analysis

Data were first epoched into trials -0.5 s pre-stimulus and 1.5 s post-stimulus onset. The data were then baselined using the 500 ms pre-stimulus time window. Semi-automated artefact rejection was completed using the Fieldtrip toolbox (Oostenveld, Fries, Maris, & Schoffelen, 2011) as well as manually inspecting each trial for muscle and electrical interference. ICA was run on the remaining data and artefact components were visually identified and removed. The average reference was then computed. A linearly constrained minimum variance (LCMV) beamformer (Van Veen, Van Drongelen, Yuchtman, & Suzuki, 1997) was applied with an 8mm resolution and a broad 30-9 0 Hz bandpass filter using the template headmodel provided with fieldtrip. The result was projected into the Montreal Neurological Institute (MNI) coordinate system. The coordinate contributing the peak gamma power intensity was selected and used to construct a virtual sensor (Figure 1b-c).

**Figure 1.**
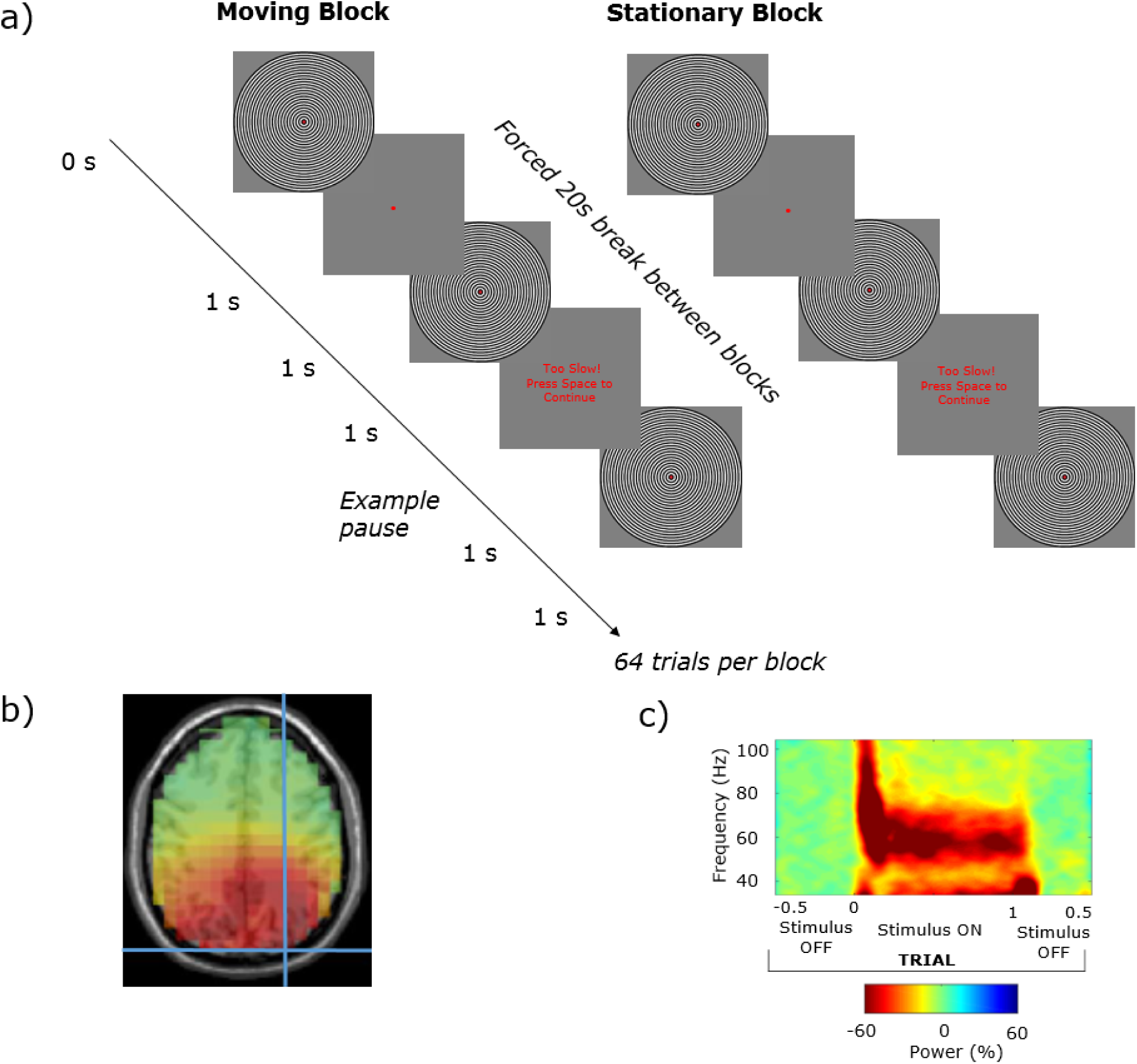
a) An annular grating was used to induce visual gamma oscillations. For each trial the stimulus was on the screen for 1s, and off for approximately 1 s. The task was a block design with 84 trials per block, the stimuli was either moving or stationary for each of the 6 blocks. Participants were required to press the spacebar when the stimuli disappeared from the screen and were prompted if their response was too slow. Following source localisation a virtual sensor of the peak gamma intensity was chosen. b) A single participants peak gamma at MNI coordinate 27 -90 36. c) The time-frequency spectrogram (obtained using Hilbert transformation of bandpass filtered data) of gamma activity at this virtual sensor in relation to trial timing, exemplifying induced visual gamma activity and how it is time-locked to the grating stimulus (in this case moving).

In order to estimate the peak frequency of gamma, a bootstrapping method was used as outlined in Magazzini et al. (2016) that produces a metric of the peak frequency reliability as a form of quality control (QC). Using this method, a spectral analysis was first completed that used a smoothed periodogram (Bloomfield, 2004), Fourier method. For each trial, the time series is demeaned and tapered with a Hanning window. The raw periodogram was then computed for the baseline (-500ms) and sustained gamma (250-750ms) time periods, and smoothed with a Gaussian kernel. The single-trial spectra was averaged across trials for the baseline and stimulus separately. From this, the amplitude spectrum was calculated as percent change from baseline. Bootstrapping was performed with 10,000 iterations on the spectral data with a frequency-window of 35-90 Hz. Trials were resampled with replacement. Peak frequency was calculated as the greatest increase from baseline of the average of the resampled single-trial data. The QC procedure provided a metric of reliability by calculating the frequency width required to accommodate 50% of the bootstrapped frequencies around the distribution mode. In this study a threshold of +/-1 Hz was applied, meaning that if more than 50% of the iterations fell outside 1 Hz either side of the mode, the dataset was rejected. Moving gratings peak frequency estimation QC passed all 18 data sets (Supporting Information Figure S1), indicating reliable gamma peaks could be estimated for all participants. Peak frequency estimation QC for the static grating task passed 9/16 data sets indicating, as expected (Muthukumaraswamy & Singh, 2013), that the static grating produces a lower signal-to-noise ratio than moving gratings (Supporting Information Figure S2).

### DCM

Dynamic Causal Modelling for Steady State Responses (DCM-SSR) was conducted using the methods of Shaw et al. (2017) which utilised a variation on the canonical microcircuit (CMC) as a neural model. DCM uses a generative model comprising the neural-mass model as well as an observation model, which when using EEG or MEG is typically a lead-field weighting (Moran et al., 2009; Shaw et al., 2017). The neural mass model for CMC comprises 4 types of interacting cell populations. For these the variation on the neural model employed by Shaw et al. (2017) includes six types of parameter, including time-constants (T), local (G) and extrinsic (A) synaptic connectivity strengths, exogenous input (C) strength, delay (D) and presynaptic firing (S). Of particular interest is the modulation of intrinsic or local connectivity (parameter G) but also population time-constants (parameter T) (Figure 2), these and an overall gain parameter (L) are allowed to vary all other parameter types outlined above are fixed in the model.

Under this model it was possible to characterise the local synaptic connectivity between inhibitory interneurons and excitatory pyramidal, and stellate cells (Figure 2). Two non-reciprocal excitatory connections exist, one between L4 stellate to L2/3 pyramidal cells, the second between L2 pyramidal to L5/6 pyramidal cells. Each population also has an inhibitory, self-modulatory gain connection modelled by G1, G4, G7, and G10. Reciprocal L2/3 pyramidal to interneuron connection are modelled by the G11 and G12 parameters that correspond physiologically to the generators of gamma rhythm, according to the PING model (Bartos et al., 2007; Tiesinga & Sejnowski, 2009; Whittington et al., 2000). Although this model may be overly-simplified in terms of the true cytoarchitecture, mean-field models balance biological complexity against computational estimability. Indeed, the model has been shown to be sufficiently detailed enough to recapitulate variations in local connectivity induced by subtle pharmacological manipulation (Shaw et al., 2017).

**Figure 2.**
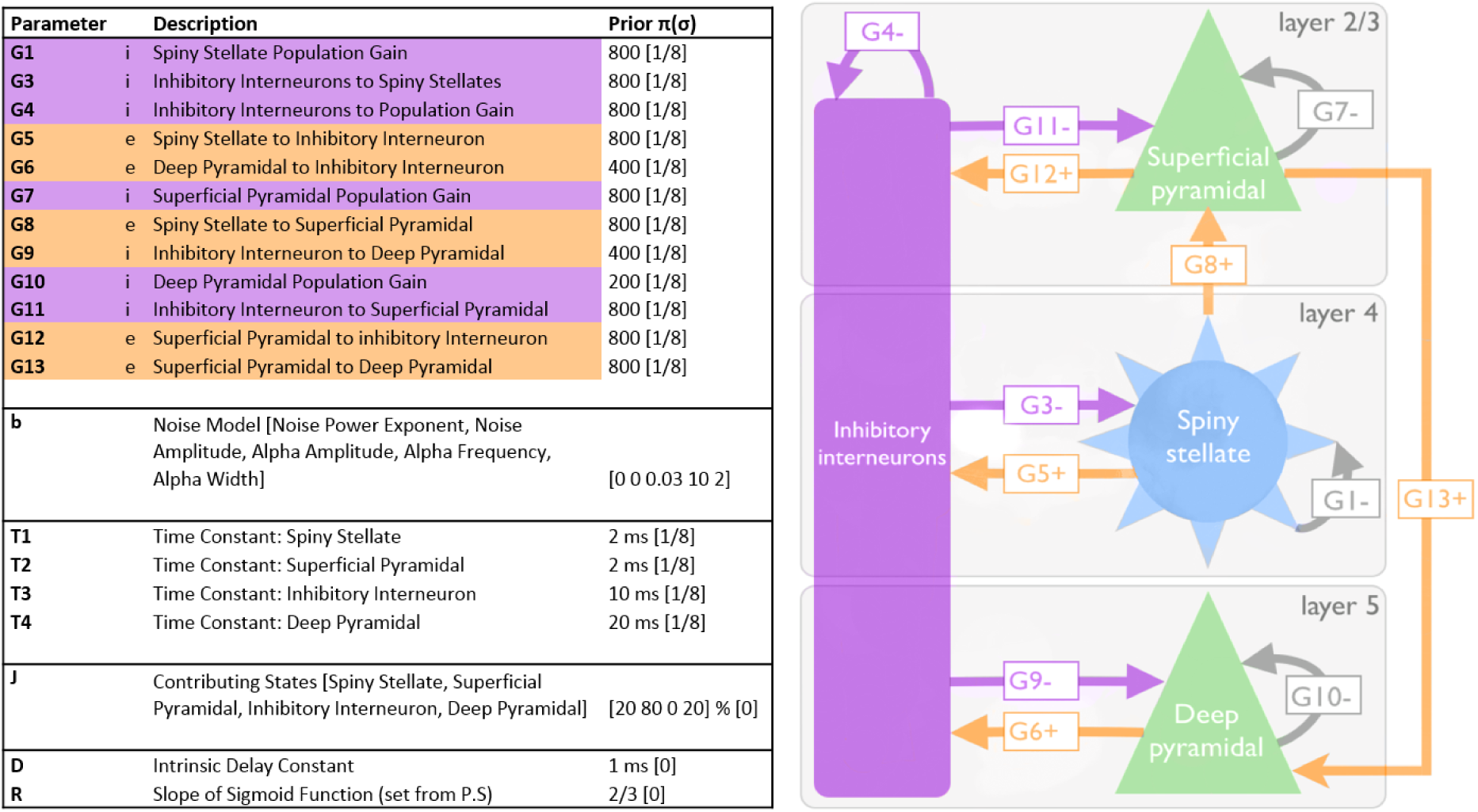
Adapted from Shaw et al. (2017). Depicts and describes the canonical microcircuit (CMC) used in the DCM procedure. *Left* Three-layer model, excitatory connections are shown in orange, and inhibitory (i.e. GABAergic) connections in purple. Grey arrows represent self-inhibition within each of the excitatory cell populations. *Right* descriptions of parameters that define the model, including their prior values and their precisions (sigma).

Initial starting values for each of the model parameters were derived by first fitting the CMC model to the mean spectral density across participants and conditions. These values form the priors that are used in the individual DCM-SSR fits for each of the individual datasets. G1, G3, G10 and G13 were fixed as in the study by Shaw et al. (2017) where these parameters were found to have little to no effect on the fitted spectral density. Likewise, we also fixed T1 as it was found to have a profound effect on model stability (Shaw et al., 2017). The output provided values of individual parameter strengths for each condition.

## Results

### Blood sampling and cycle timing confirmation

All participants were confirmed to have progesterone and oestradiol levels in the range consistent with the follicular and luteal phases on the day they participated in each study session (Figure 3)

**Figure 3.**
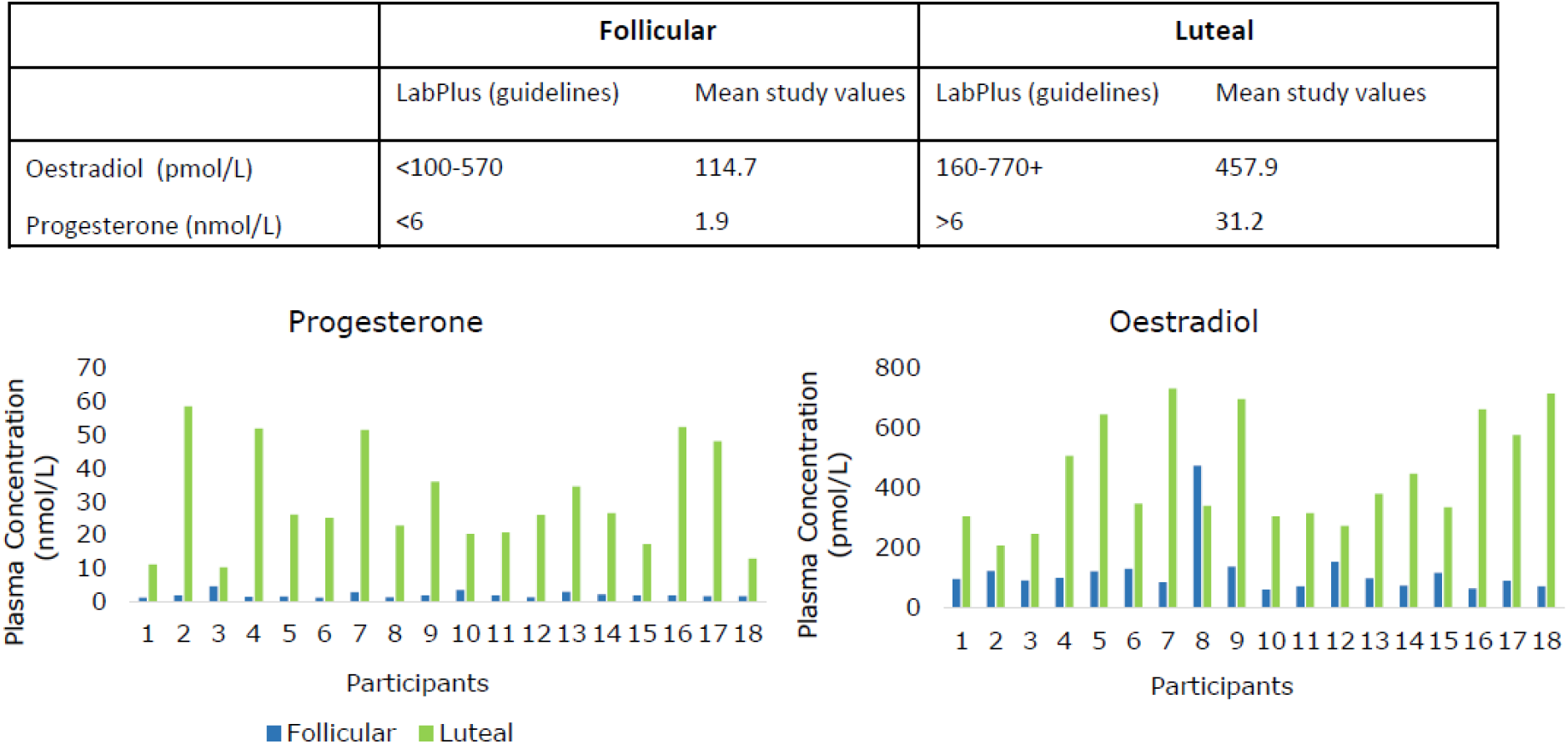
Plasma levels of progesterone and oestradiol taken at the EEG recording to confirm cycle timing. The data indicates that 100% of participants were in the correct phase at the time of testing.

Moving gratings peak frequency estimation QC passed all 18 data sets meaning all participants were included in the subsequent analyses. All Wilcoxon-signed rank tests were subjected to False Discovery Rate (FDR) correction for multiple comparisons (Benjamini & Hochberg, 1995). A Wilcoxon-signed rank test showed that participants had significantly higher peak mean gamma frequency in the luteal phase (M = 63.42 Hz, SD = 5.30 Hz) compared to the follicular phase (M = 59.86 Hz, SD = 7.19 Hz; Z = -2.42, p = 0.032 FDR) (3.56 Hz difference) (Figure 4). The corresponding peak amplitudes were also subjected to a Wilcoxon signed-rank test. However there was no significant difference in percent signal change between peak amplitude for the luteal (M = 111.01 %, SD = 66.51Hz) and follicular phases (M = 95.72 %, SD = 44.70 %; Z = -0.21, *p* = 0.836 FDR) (Figure 4).

**Figure 4.**
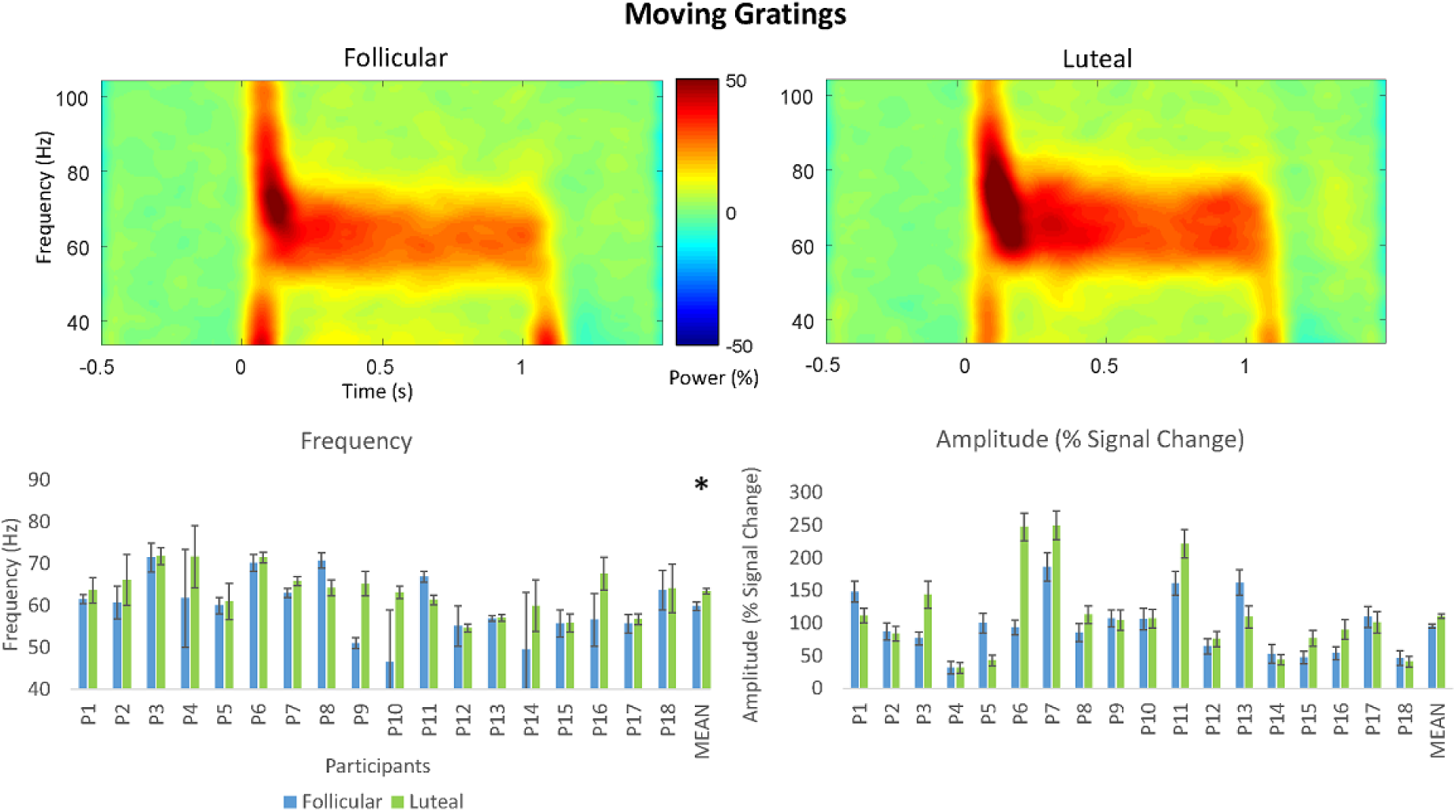
*Top* Grand-averaged time-frequency spectrogram for the follicular and luteal phases for the moving grating stimulus type. Intensity of colour warmth indicates changes in power (%) from the baseline. *Bottom* individual participant frequency and amplitude results. Study means show significantly higher peak mean frequency in the luteal compared to follicular phase. No significant differences were found for amplitude. Error bars show standard deviation of the bootstrapped distribution.

Peak frequency estimation QC for the static grating task passed 9/16 data sets to be included in subsequent analyses. As mention above, this is expected [44], as the static grating produces a lower signal-to-noise ratio than moving gratings (Supporting Information Figure S2). Despite the low numbers, consistent with the finding for moving gratings, a Wilcoxon-signed rank test also showed that participants had significantly higher peak mean gamma frequency in the luteal phase (M = 58.16 Hz, SD = 3.95 Hz) compared to the follicular phase (M = 52.41 Hz, SD = 3.00 Hz; Z = -2.43, *p* = 0.032 FDR) (5.75 Hz difference) (see Figure 5). There was also no significant difference in percent signal change of peak mean amplitude for the luteal (M = 114.49 %, SD = 57.12 %) compared to the follicular phase (M = 120.84 %, SD = 71.19 %; Z = -0.42, p = 0.836 FDR) (Figure 5).

**Figure 5.**
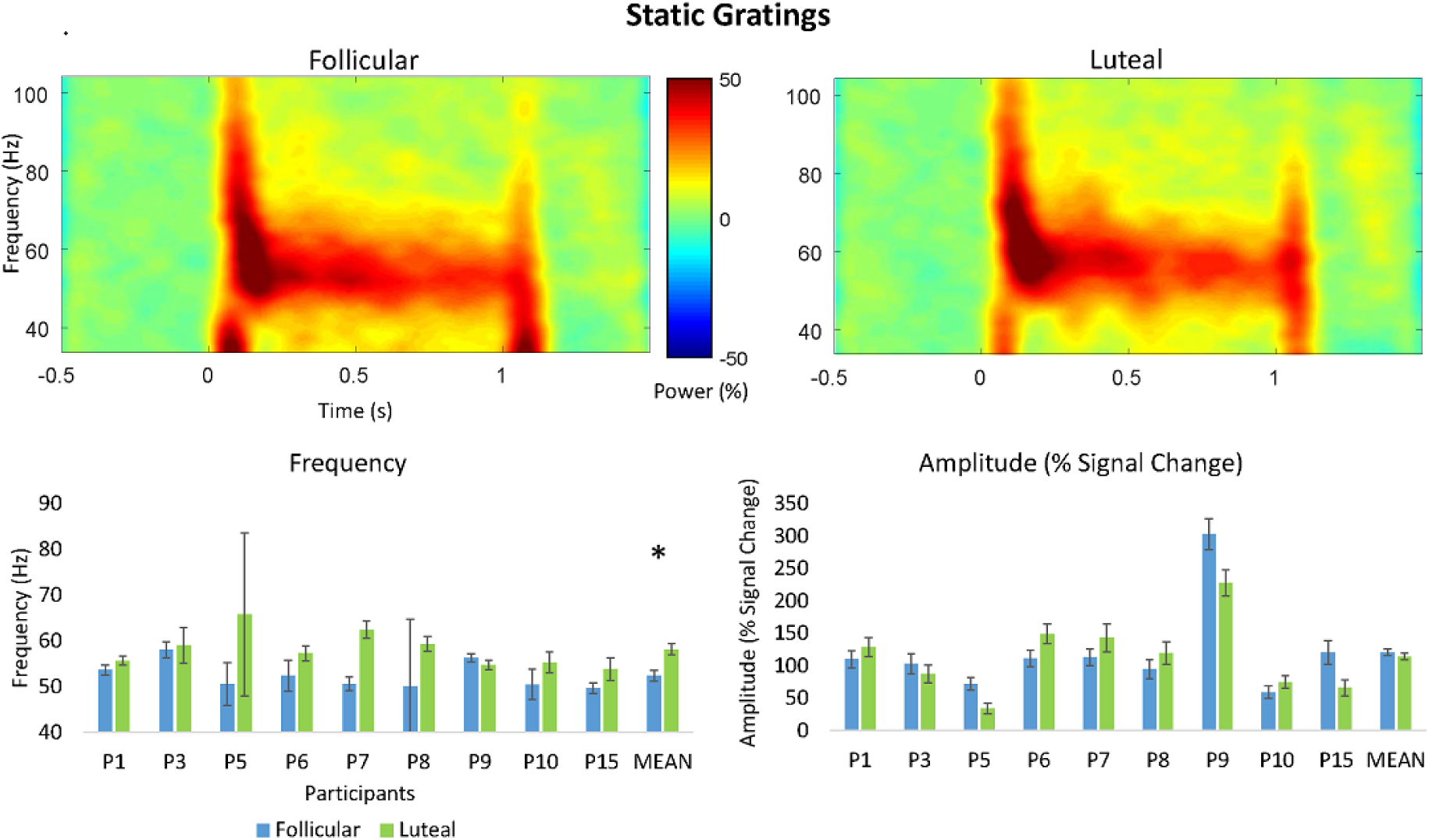
*Top* Grand-averaged time-frequency spectrogram for the follicular and luteal phases for the moving grating stimulus type. Intensity of colour warmth indicates changes in power (%) from the baseline. *Bottom* individual participant frequency and amplitude results. Study means show significantly higher peak mean frequency in the luteal compared to follicular phase. No significant differences were found for amplitude. Error bars show standard deviation of the bootstrapped distribution.

Spearman’s correlations were conducted to check for potential correlations between change in amplitude between the luteal and follicular phase, change in frequency and oestradiol or progesterone (Figure 9). For moving gratings no significant correlation was found between progesterone (M = 28.66 nmol/L, SD = 15.76 nmol/L) and frequency (M = 3.57 Hz, SD = 6.38 Hz; rs =0.388, p = 0.111) or percent signal change of peak mean amplitude (M = 15.29 %, SD = 49.94 %; rs = -0.120, p = 0.636). Likewise, no significant correlation was found between oestradiol (M = 326.61 pmol/L, SD = 217.28 pmol/L) and frequency (rs = 0.442, p = 0.066) or percent signal change of peak mean amplitude (rs = -0.178, p = 0.481). Similarly, for static gratings, no significant correlation was found between progesterone (M = 25.56 nmol/L, SD = 15.02 nmol/L) and frequency (M = 5.76 Hz, SD = 5.40 Hz; rs = 0.452, p = 0.222) or percent signal change of peak mean amplitude (M = -6.34 %, SD = 41.04 %; rs = 0.218, p = 0.574). Likewise, no significant correlation was found between oestradiol (M = 354.22 pmol/L, SD = 198.79 pmol/L) and frequency (rs = 0.617, p = 0.077) percent signal change of peak mean amplitude (rs = 0.383, p = 0.308).

## Dynamic Causal Modelling

### Cycle and mean parameter strength

A repeated measures ANOVA was run to explore the effects of menstrual cycle phase (follicular versus luteal), and grating (moving versus static) on the G (local connection), and T (time constant) parameters (Figure 6). There was no significant phase by grating-type interaction (F_(1,10)_ = 0.38, *p* = 0.866). However, because the primary purpose of this analysis is to explore the effect of menstrual cycle phase on the parameters, the main effects are interpreted. A significant main effect of menstrual cycle phase was found (F_(1,10)_ = 4465.82, *p* = 0.012). Two parameters showed this main effect of phase. G7 (F_(1,10)_ = 6.77, *p* = 0.026), and G9 (F_(1,10)_ = 7.10, *p* = 0.024). Comparison of estimated marginal means shows that G7 is stronger in the luteal phase (M = 1.33, SE = 0.056) than the follicular phase (M = 1.18, SE = 0.044). G9 is stronger in the follicular (M = 2.11, SE = 0.028) than the luteal phase (M = 2.03, SE = 0.029). No significant main effect of grating was found (F_(1,10)_ = 34.88, *p* = 0.131).

**Figure 6.**
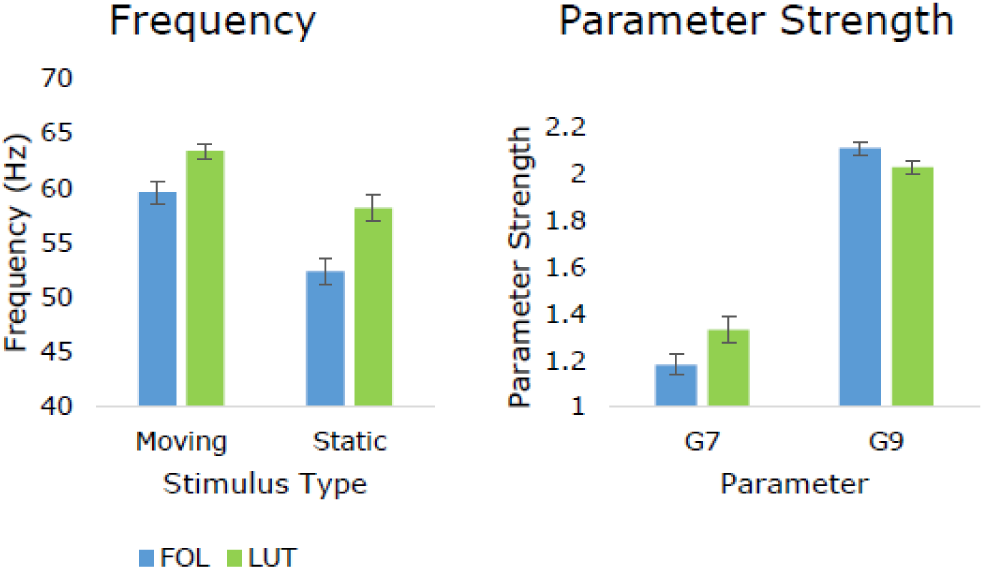
*Left* peak mean frequencies from the QC estimation show significantly greater frequency in the luteal compared to follicular phase for both moving (+3.56 Hz) and static gratings (+5.75Hz). These are shown side-by-side with *Right* the significant difference in estimated marginal means of the parameter strengths for G7 and G9 in the follicular compared to luteal phase. G7 is significantly stronger in the luteal phase, G9 is significantly stronger in the follicular phase. Error bars show standard error.

## Discussion

Using EEG to record visual gamma oscillations during the luteal and follicular phases of the menstrual cycle, this study found significantly higher peak gamma frequency in the luteal phase compared to the follicular phase. This result shows that endogenous modulation of gamma oscillations can be reliably measured using EEG. Evidence of this ability is important as the majority of existing work with gamma oscillations has been completed using MEG. This is largely due to the relatively poorer signal-to noise ratio of EEG and historical lack of advanced signal processing techniques to overcome this. However, EEG is both cheaper and more widely available both at academic institutions and hospitals than MEG, an important factor for future applications of this research.

Making any detailed inferences about specific neurobiological causes for our findings are complicated and limited by the observational nature of the study design. However, the lack of any correlation between peripheral measures of progesterone or oestradiol and peak gamma amplitude or frequency do not support the argument that absolute neurosteroid concentrations are the key contributors to the effects of menstrual cycle found in this study, as peripheral concentrations of progesterone and oestradiol have been found to be well correlated with concentrations in the brain (Bixo, Bäckström, Winblad, & Andersson, 1995; Wang, Seippel, Purdy, & Bäckström, 1996). Although we did not directly measure allopregnanolone levels, these are correlated with progesterone, particularly in the luteal phase (Wang et al., 1996). Instead, because gamma oscillations in V1 have been correlated with GABA_A_ receptor properties such as density in human visual cortex (Kujala et al., 2015) this suggests that receptor dynamics may be more important.

The pharmaco-MEG literature provides a number of examples showing that increasing GABAergic inhibition via GABA enhancing drugs leads to a decrease in gamma frequency (Campbell et al., 2014; Lozano-Soldevilla et al., 2014; Magazzini et al., 2016). In light of this our finding of relatively higher frequency in the luteal compared to the follicular phase found in this study could be interpreted as greater GABAergic inhibition in the follicular phase. This is biologically feasible if mechanisms for allopregnanolone tolerance are the primary mediator of the balance of inhibition during the mid-luteal phase, as during the luteal phase allopregnanolone is at its highest concentration in the brain. Based on animal literature the primary mediator of apparent changes in GABAergic inhibition is related to upregulated expression of GABA_A_ receptors containing α4 and δ subunits (Smith et al., 2007). Therefore a potential decrease in sensitivity may represent a break down or modulation of α4 containing receptors found in mice (Lovick et al., 2005) occurring in the late luteal phase of the human female menstrual cycle. This may also account for the decrease in sensitivity to administered allopregnanolone found in the study by Timby et al. (2016) If the mechanism for reported changes in GABAergic inhibition are related to α4 receptors, this may be accounting for the conflict between the findings with administered benzodiazepines and allopregnanolone (Sundstrom et al., 1997; Timby et al., 2016). Decreased sensitivity to benzodiazepines and relatively unchanged or increased sensitivity to endogenous neurosteroids such as allopregnanolone is associated with increases in α4 containing GABA_A_ receptors (Maguire et al., 2005; Wafford et al., 1996), though it is less clear why the findings for pregnanolone and allopregnanolone are contrasting (Sundstrom et al., 1998). However, allopregnanolone is a far more potent modulator of GABA than pregnanolone (Zhu, Wang, Bäckström, & Wahlström, 2001). It has been proposed that tolerance may be different depending on the potency of the modulating neurosteroid (Timby et al., 2016).

In this study we only found a change in gamma frequency and not amplitude. This has been found in one other study on the effects of tiagabine on visual gamma oscillations (Magazzini et al., 2016). In our case, this may have been due to EEG signal to noise ratio being generally lower than that of MEG (Muthukumaraswamy & Singh, 2013). However, this is unlikely as the reliability of the peak frequency estimation was as good as, or at least comparable to, the findings of Magazzini et al. (2016), indicating high signal-to-noise ratio. It has also been proposed that the neural mechanisms behind amplitude and frequency of gamma can be differentially modulated, with frequency more linked to the time-constant of inhibitory processes (Magazzini et al., 2016). To support this, when the data for the tiagabine data as presented by Magazzini et al. (2016) was subjected to DCM-SSR by Shaw et al. (2017), it was found that individual variability in the time constant of inhibitory interneurons was found to be significantly modulated by gamma frequency but not amplitude. Furthermore, a contribution analysis was completed to determine the key parameters contributing to gamma frequency. This was found to be parameter G7; the self-inhibition of superficial pyramidal cells. By contrast, gamma amplitude was found to be primarily determined by G11; the strength of the inhibitory interneuron to superficial pyramidal cell connection.

In the current DCM analyses we found several microcircuit parameters that were modulated by menstrual cycle phase; parameters G7 and G9. As explained above, G7 represents superficial pyramidal self-inhibition and was increased in the luteal phase. Whereas G9 represents inhibitory interneuron connections to deep pyramidal cells and was increased in the follicular phase. The previous contribution analysis run by Shaw et al. (2017) determined four parameters with the greatest contribution to beta and gamma peak amplitude and frequency by testing a parameter’s sensitivity to variation in each respective spectral feature, two of which are relevant here. Parameter G7 (self-inhibition of superficial pyramidal cells) was the predominant determinant of peak gamma frequency where increasing G7 leads to corresponding increases in gamma frequency. This is entirely consistent with the current results. From a neurobiological perspective, G7 as a self-inhibition parameter is a “lumped” parameter that could subsume a variety of potential gain-control mechanisms, including neuromodulation, receptor cycling or receptor desensitisation, any of which could be at play across the endogenous menstrual cycle. Furthermore, any of these mechanisms may feasibly be modulated by GABAergic changes. As has already been mentioned, in response to increases in allopregnanolone, the primary mediator of apparent changes in inhibition is upregulated expression of GABA_A_ receptors containing α4 and δ subunits (Smith et al., 2007). δ-GABA_A_ receptors are extrasynaptic mediators of tonic inhibition and have been related to gain-control mechanisms (Semyanov, Walker, Kullmann, & Silver, 2004; Stell, Brickley, Tang, Farrant, & Mody, 2003). They are also at particularly high density in layer 2/3 (Drasbek & Jensen, 2006), where G7 is modelled.

With respect to G9 representing inhibitory interneuron to deep pyramidal cells, Shaw et al. (2017) found this parameter was positively correlated with beta rather than gamma amplitude. G9 more directly suggests modification of GABAergic interneuron system, although specifically in its laminar interaction with the deep pyramidal cells. Interestingly, and in contrast to the above, this modification indicates that there is greater inhibition in the follicular phase than luteal phase. This is proposed to be in keeping with one of the unique actions of allopregnanolone at endogenous levels. Below certain levels allopregnanolone can produce a paradoxical effect and depress inhibition via polarity dependent action on Cl^-^ influx and efflux (Bäckström et al., 2011; Smith et al., 2007). However, δ-GABA_A_ receptors are more sensitive to lower levels of allopregnanolone, and in the healthy menstrual cycle, still produce an overall increase in GABAergic inhibition when allopregnanolone levels increase endogenously (Belelli et al., 2009; Lovick et al., 2005; Maguire et al., 2005; Smith et al., 2007). Research has shown that there is reduced expression of δ-GABA_A_ receptors in layer 5 (Drasbek & Jensen, 2006) where parameter G9 is modelled. Rather, tonic inhibition is modulated by α5-GABA receptors which allopregananolone has a much lower affinity for (Peng et al., 2009) but can also produce a paradoxical effect (Burgard, Tietz, Neelands, & Macdonald, 1996; Smith et al., 2007). The reduced sensitivity to allopregnanolone within layer 5 due to the absence of δ-GABA_A_ receptors may be leading to the opposing difference in inhibition found in the luteal compared to the follicular phase.

Taken together, these modifications suggest a change in the laminar functioning of the (visual) cortex across the menstrual cycle with relatively less superficial pyramidal cell activity in the luteal phase and relatively more GABAergic inhibition of deep-level activity in the follicular phase. In invasive animal recordings, gamma rhythm activity predominates in the superficial layers whereas in deeper layers beta oscillations are more dominant (Maier, Adams, Aura, & Leopold, 2010; Xing, Yeh, Burns, & Shapley, 2012). From a theoretical perspective, this may suggest alterations in hierarchical prediction coding mechanisms where it has been suggested that superficial cells encode ascending prediction errors while predictions are encoded by deep pyramidal cells and then transmitted to lower levels of the cortical hierarchy (Bastos et al., 2012; Friston, Bastos, Pinotsis, & Litvak, 2015). On the basis of the present data alone, any argument we could make about prediction coding mechanisms across the menstrual cycle would be speculative, but we note that in the same experimental cohort, we recorded not only resting-state EEG but mismatch negativity (MMN) data. The MMN in particular specifically allows measures of hierarchical predictive coding to be made - albeit typically in the auditory system (Garrido, Kilner, Kiebel, et al., 2009; Garrido, Kilner, Stephan, & Friston, 2009). As such, although it is beyond the scope of the current work, it is possible that in the near future we may be able to explicitly test these speculations regarding predictive coding.

As well as providing a valuable tool for measuring functional changes in the brain related to changes in balance of excitation and inhibition, one of the most important impacts of our research is that it shows that endogenous changes across the menstrual cycle significantly affect the EEG signal during visual gamma tasks. Our study also makes clear that consideration of menstrual timing in women is important if they are to be included in either healthy control or patient studies. This is particularly relevant for repeated-measures data where recordings may take place at different phases of the menstrual cycle. One way to overcome this would be to ensure participants all came in on the same phase of their menstrual cycle. The follicular phase is by far the most straightforward to verify as its initiation is signalled by the onset of menstrual bleeding. Alternatively, females on oral contraception, whilst taking active hormone pills will be at a predictable and constant stage in their cycle. Any other form of hormonal contraception or therapy should also have its specific impact on the menstrual cycle and gonadal hormones taken into account. For example modern contraceptive implants, such as *Implanon,* can lead to increased irregularity of menstrual bleeding so in these cases follicular phase would be again easier to estimate (Mansour, Korver, Marintcheva-Petrova, & Fraser, 2008).

### Strengths, limitations and future directions

A key strength of this study was the rigorous cycle tracking and confirmation of cycle timing. Participants were tracked for three full cycles prior to their first study date. Participants with any irregular cycle lengths, despite often self-reporting regular cycles, came in the day before a potential study date for a blood sample. The study date was rescheduled if they were not in their luteal phase. This happened on a small number of occasions. Within the average menstrual cycle there is very common, and healthy, variance in intra-individual cycle length and regularity (Fehring, Schneider, & Raviele, 2006). This leads us to the conclusion that in studies without blood or saliva quantification of progesterone, it cannot be unequivocally stated that testing took place during the luteal phase. Even counting back from menstrual onset after the study, though successful in some cases, will not pick up on anovulatory cycles. Anovulatory cycles have no surge in hormones during the luteal phase and can affect up to 38% of women aged 20-24 for at least 1/3 cycles (Metcalf & Mackenzie, 1980). In addition, as already mentioned, all study sessions began between 2-4pm to control for diurnal variations in neurosteroid levels (Tiihonen Möller et al., 2016).

One of the limitations of this study was the apparent trade-off between moving and static gratings. Moving gratings produced the cleanest data according to the QC thresholding. This lead to the greatest number of useable datasets to take forward through data processing. It has been reported previously that visually induced gamma oscillation can hit a frequency ceiling at around 70 Hz and that moving gratings tend to induce gamma at a frequency closer to 70 Hz than static (Swettenham, Muthukumaraswamy, & Singh, 2009). This was referred to as a potential saturation of the visually induced gamma effect (Swettenham et al., 2009). The relatively smaller mean change in gamma frequency between cycle phases for moving (3.56 Hz) compared to static gratings (5.75 Hz) may be some indication of this occurring in our data. A potential criticism of our data analysis may come from the nature of the QC approach to analysing visual gamma; where data that does not contain robust gamma gets rejected. This means that individuals that do not produce or record robust gamma get rejected alongside poor quality datasets. This raises questions around whether results are representative of the general population, an important point also echoed by Magazzini et al. (2016). However all of our participants passed QC for at least the moving stimulus so this effect appears to be representative for the majority of our participants and therefore potentially the larger population of at least young women.

To explore the apparent changes in the GABA system further in the future, it would be useful to record induced gamma oscillations at the very beginning of the luteal phase as well as the end to compare changes over time. This may help disentangle and explain some of the contradictory findings in the human literature as well as provide a dimension of long-term changes that are not provided in studies of the far shorter menstrual cycle of rodents (4 days). In addition, to exclude a potential interaction of oestradiol, a recording during ovulation when progesterone levels are low and oestradiol high may provide important insight, especially as oestradiol has a depressive effect on GABAergic inhibitory input (Wójtowicz & Mozrzymas, 2010). However, it should be noted this was not considered to be a major limitation to our inferences regarding allopregnanolone as oestradiol has been shown to be less influential on overall GABA levels, and hence inhibition, than the effect of progesterone and its metabolites in humans (Epperson, Haga, Mason, & et al., 2002; Epperson et al., 2005). Indeed, it is progesterone and it’s metabolites that have a more clearly defined implication in the GABA-linked menstrual disorders such as PMDD (Bäckström et al., 2014; Barth, Villringer, & Sacher, 2015; Epperson, Haga, Mason, & et al., 2002; Girdler, Straneva, Light, Pedersen, & Morrow, 2001; Reddy, 2004). Furthermore, in terms of the value of such an approach, the peak during ovulation is far shorter than that of the luteal phase and may not capture the longer term dynamics, instead offering an entirely unique piece of information. As such, it would be difficult, if not impossible, to quantify the contribution of progesterone and oestradiol independently in studies of purely endogenous changes in the mid-late luteal phase.

In conclusion, this study provides evidence for menstrual-cycle related changes in visual gamma oscillations. Increased gamma frequency was found in the luteal compared to the follicular phase, thus demonstrating endogenous synaptic modification of excitation-inhibition occurring across the menstrual cycle. Our findings also indicate that there are complex functional changes within the cortical microcircuitry across the menstrual cycle and exemplify the potential value of DCM in elucidating the mechanisms behind these changes, in addition to analyses of spectral data features.

## Acknowledgements

RLS is supported by Auckland Medical Research Foundation Doctoral Scholarship. SDM is supported by a Rutherford Discovery Fellowship administered by the Royal Society of New Zealand. AS is supported by a Wellcome Trust Strategic Award (104943/Z/14/Z). KDS is supported by the UK MEG Partnership Grant (MRC/EPSRC, MR/K005464/1), CUBRIC and the School of Psychology at Cardiff University. The work was supported by a Faculty Research Development Fund grant.

The authors have no conflicts of interest to declare.

## Author contributions

Conceptualisation, RLS, and SDM; Methodology, RLS, SDM, ADS, KDS; Investigation RLS, SDM, RJM; Formal Analysis, RLS; Writing – Original Draft, RLS and SDM. Writing – Review and Editing, RLS, SDM, ADS, KDS, RJM, FS. Funding Acquisition, SDM. Supervision, SDM, FS.

